# Quantitative interactions drive *Botrytis cinerea* disease outcome across the plant kingdom

**DOI:** 10.1101/507491

**Authors:** Celine Caseys, Gongjun Shi, Nicole Soltis, Raoni Gwinner, Jason Corwin, Susanna Atwell, Daniel Kliebenstein

**Author notes:** **Correspondence:** Daniel J. Kliebenstein, Department of Plant Sciences, University of California, Davis, One Shields Ave, Davis, CA, 95616, USA.

## Abstract

*Botrytis cinerea* is a polyphagous fungal pathogen that causes necrotic disease on more than a thousand known hosts widely spread across the plant kingdom. While it is known that quantitative resistance in the host and quantitative virulence in the pathogen largely mediate this pathosystem, how this pathogen interacts with the extensive host diversity is unknown. Does this pathogen have quantitative virulence efficiency on all hosts or individual solutions for each host? To address this question, we generated an infectivity matrix of 98 strains of *Botrytis cinerea* on 90 genotypes representing eight host plants. This experimental infectivity matrix showed that the predominant sources of quantitative variation are between host species and among pathogen strains. Furthermore, the eight eudicot hosts interacted individually with *Botrytis cinerea* strains independently of the evolutionary relatedness between hosts. An additive quantitative model can explain the complexity of these interactions in which Botrytis host specificity and general virulence have distinct polygenic architectures.

## Introduction

Plants live in complex ecosystems and are under constant combinatorial abiotic and biotic pressure. Within biotic interactions, a plant can be exposed to multiple pathogens with various attack strategies. As counter-measures to the diversity of pathogen attacks, plants have developed multi-layered resistance mechanisms ranging from constitutive physical defenses to inducible responses coordinated by the plant immune system upon the recognition of danger (Glazebrook, 2005, Herman and Williams, 2012, Wilkinson et al., 2019). While all plants have a multi-layered defense system, the individual components have complex and diverse evolutionary histories. Defense components such as resistance genes (Jacob et al., 2013) or specialized defense compounds (Chae et al., 2014) are frequently specific to limited lineages or even individual plant species. In contrast, other components such as the cell wall (Sørensen et al., 2010), defense hormone signaling (Berens et al., 2017) or reactive oxygen species (Inupakutika et al., 2016) are widely shared across plant lineages. It is not fully clear how the various histories of these defense components differentially contribute to the interaction with pathogens across plant species.

To counter the variety of plant defenses, pathogens use diverse virulence mechanisms to attack and/or interfere with the host resistance. The genetic architecture of these virulence mechanisms varies across the continuum of different pathogens lifestyles and host specificity (Barrett et al., 2009). At one end of the continuum, specialist biotrophs develop intricate relationships, living and feeding within the host tissues. At the other end, generalist necrotrophs attack, kill and feed on degraded tissues (Möller and Stukenbrock, 2017). Host specificity also covers a large range from specialists attacking one or few hosts to generalists that can have multiple hosts, taxonomically related or not. Both lifestyle and host-specificity can influence the genetic architecture of the plant-pathogen interaction (Morris and Moury, 2019). Biotrophs and specialist pathogens tend to have a narrow co-evolution with their host that generates a qualitative genetic architecture dominated by a few interaction-specific large-effect genes in both host and pathogen. At the opposite, most generalist interactions rely on quantitative genetic architectures with small effect loci (Corwin and Kliebenstein, 2017, Poland et al., 2009, Lannou, 2012).

In this study, we focus on a panglobal and ubiquitous necrotroph, *Botrytis cinerea* (grey mold; Botrytis hereafter) to estimate the relative contribution of host genetic variations, from wild and domesticated genotypes to species from different clade, in shaping the interactions with a fungal pathogen. Botrytis causes billions/annum of crop damage both pre- and post-harvest to various crops from ornamentals to vegetables and fruits (Veloso and van Kan, 2018, Fillinger and Elad, 2015). It has a broad host range ranging from mosses to gymnosperms to eudicots, and is classically considered as a generalist fungal pathogen (Van Kan et al., 2017, Veloso and van Kan, 2018, Fillinger and Elad, 2015). Within the eudicots, disease symptoms are noted on 996 species widely spread across orders with no apparent phylogenetic signal or correlation with the species diversity (Fig.1). In addition to a broad host range, Botrytis also displays a polyphagous ability to infect diverse tissues within a host, such as leaf, stem or flower (Fillinger and Elad, 2015).

**Fig. 1:**
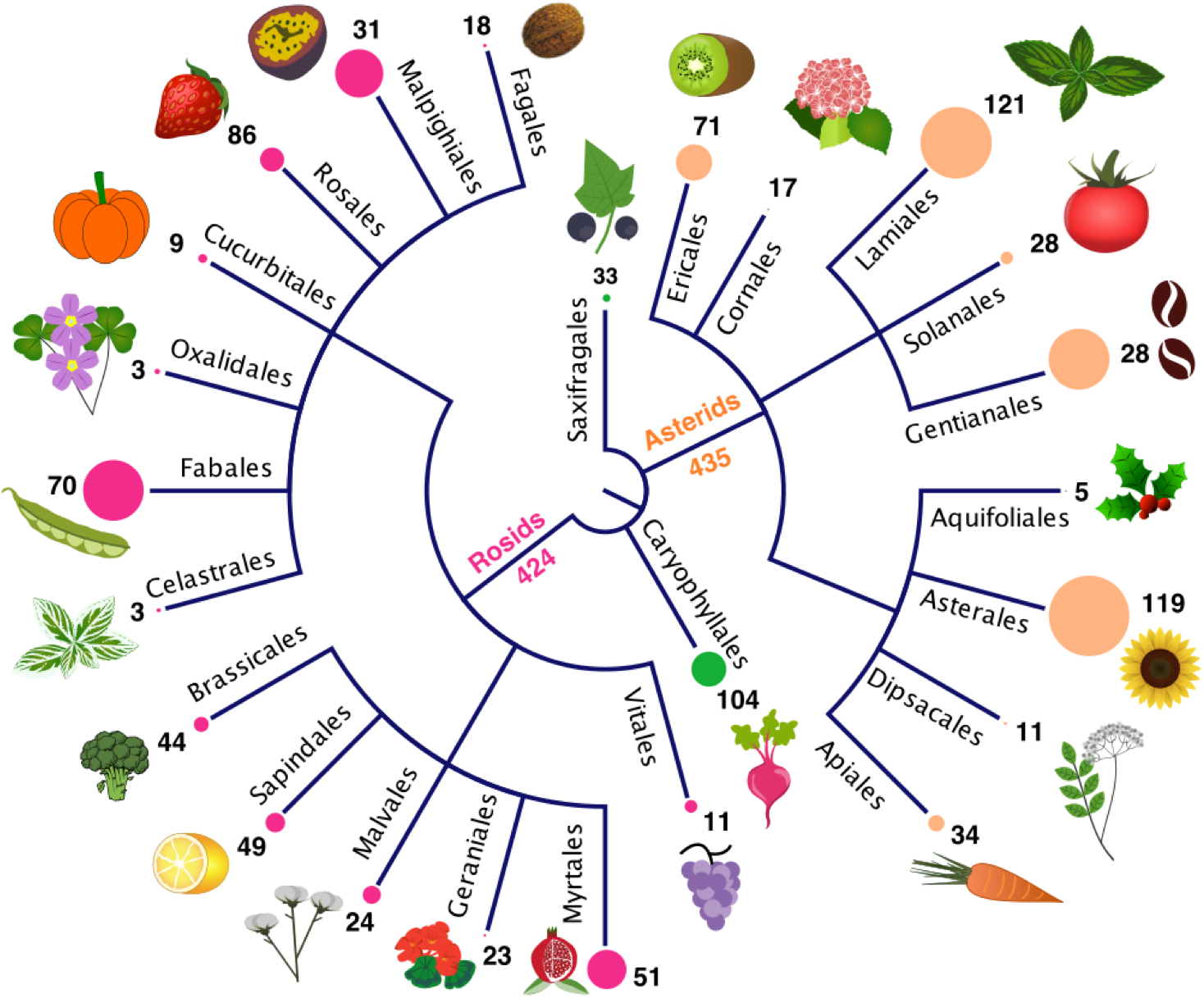
Disease symptoms caused by *Botrytis cinerea observed on plants from* the core Eudicots. Plant species with disease symptoms were extracted from (Elad et al., 2016). The tree represents major orders from the basal Eudicots (in green), Asterids (in orange) and Rosids (in pink). The size of the circle at the end of the branches is proportional to the species diversity of the order. For each order, the number of species with known disease symptom is indicated.

This ability of Botrytis to infect widely across the plant kingdom has been postulated to arise from its extensive genetic and phenotypic diversity (Atwell et al., 2015, Amselem et al., 2011, Zhang et al., 2010, Ma and Michailides, 2005, Valero-Jimenez et al., 2019, Walker et al., 2014b). Botrytis spores are widespread within the environment ranging from presence on plants to presence on non-plant substrates including within rain, indicating an ability to spread rapidly and widely (Bardin et al., 2018, Leyronas et al., 2015). This agrees with the presence of large standing genetic variation in both local and global Botrytis populations (Atwell et al., 2015, Atwell et al., 2018, Bardin et al., 2018, Calpas et al., 2006). Further increasing this diversity, the standing genetic variation is reshuffled by recombination during cold and wet months (Leyronas et al., 2015, Cosseboom et al., 2019, Walker et al., 2014b). Together, this creates a pan-global population with high levels of admixture showing limited host-specialization or local population structure. In California, where a large part of strains for this study were isolated in open fields, no evidence of host-specialization or local population structure has been found (Ma and Michailides, 2005, Cosseboom et al., 2019, Saito and Xiao, 2018, Soltis et al., 2019). In comparison, Botrytis strains obtained from closed environments like intensive greenhouse systems can display a reduced genetic diversity associated with an increase in clonal distribution (Walker et al., 2014a, Mercier et al., 2019, Diao et al., 2019). In these closed environments, Botrytis strains might specialize but it remains unclear if these events constitute examples of host plant adaptation, growth facility adaptation or genetic bottlenecks (Calpas et al., 2006, Leyronas et al., 2014, Decognet et al., 2009).

Although Botrytis strains have been isolated on a multitude of hosts, how the fungal genetic diversity interacts with the genetic diversity across plant lineages remains largely unmeasured. Because Botrytis strains infect a large array of species (Fig. 1), it is possible to empirically assess how host resistance and strain virulence contribute to the disease outcome. We generated infectivity matrices for 90 plant genotypes from eight eudicot species against 98 diverse Botrytis strains under experimental conditions in a standardized environment. This experiment aims to parse the effects of genetic diversity among and within eight hosts in interaction with Botrytis genetic diversity. In addition, we estimate the specificity and virulence of each Botrytis strain in this experimental design and their genetic architecture.

## Results

### Detached leaf assay based infectivity matrix

We focused our experiment on the eudicots (Fig. 1) that contains 71% of the plant species with noted Botrytis disease symptoms (Elad et al., 2016). Seven crop species (*Solanum lycopersicum*, *Helianthus annuus*, *Lactuca sativa*, *Cichorium endivia*, *Cichorium intybus*, *Glycine max*) were selected from amongst the known Botrytis hosts. These species cover various phylogenetic distances, centered on the Asterales, a family with large species diversity and frequently hosting Botrytis (Fig.1). Within each plant species, six genotypes were chosen to represent the genetic diversity in the high improvement germplasm (cultivar, inbred lines) while six genotypes represented the low improvement germplasm (wild, landraces) (Fig.2, Fig.S1). The crop genotypes also cover diverse geographical origins (Fig. S2, Table S1). In addition, wild type and five mutant genotypes of *Arabidopsis thaliana* were chosen to test single-layer alteration in plant resistance. To quantify disease symptoms across the eight species, we performed a detached leaf assay using a randomized replicated design. All 90 genotypes were infected with 98 *Botrytis cinerea* strains across two independent experiments to provide independent six-fold replication on each Botrytis strain x Host genotype combination (Fig. S1). In total, the infectivity matrix comprises 51,920 independent lesion measurements.

**Fig. 2:**
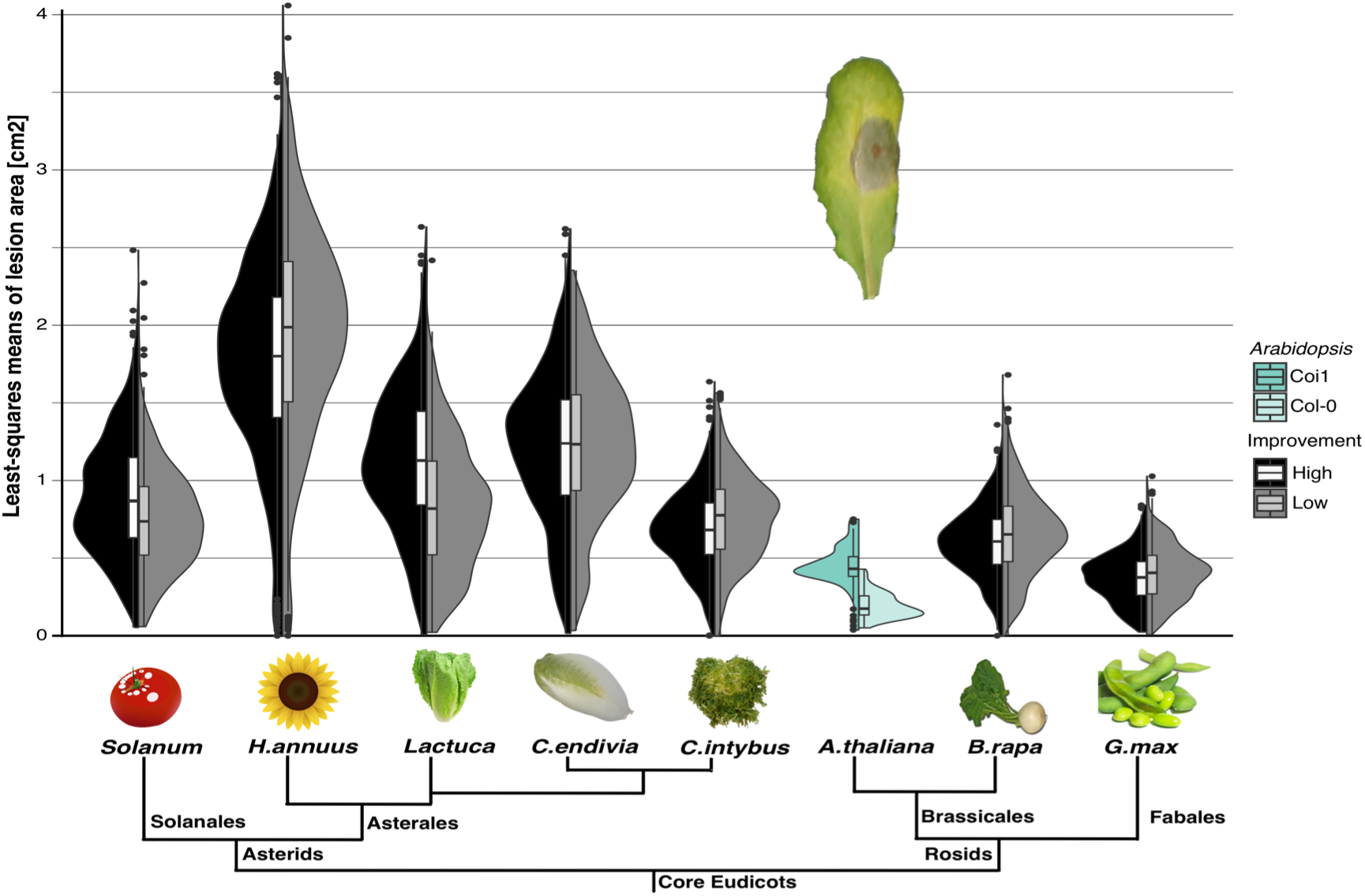
Lesion areas at 72 hours post inoculation on seven crop species shows small and inconsistent effect of domestication on the *Botrytis cinerea* interaction. Half-violins and boxplots (median and interquartile range) represent the mean lesion area distribution for genotypes with high (black, n=588) and low (grey, n=588) levels of improvement for all 98 Botrytis strains. *Lactuca* refers to *Lactuca sativa* for high improvement and *Lactuca serriola* for low improvement genotypes. *Solanum* refers to *Solanum lycopersicum* for high improvement and *Solanum pimpenellifolium for* low improvement genotypes. As reference, virulence on *A. thaliana* wild-type (Col-0, n=98) and jasmonic acid signaling mutant (*coi1*, n=98) are presented. The non-scaled tree represents the phylogenetic relationship between eudicot species.

As a common measure of the host-Botrytis interaction, we utilized lesion area on detached adult leaves. Leaves were utilized to standardize the tissue across the diverse species. Lesion area at 72 hours post inoculation (hpi) was chosen as it is an inclusive estimate that describes broad-sense heritability (e.g. genetic factors explain 89% of lesion area; Fig. 3A) from both plant and Botrytis genetic variance (Corwin et al., 2016a, Zhang et al., 2017, Soltis et al., 2019). In previous studies, we have shown that lesion area on Arabidopsis and *B. rapa* leaves is directly correlated with the area occupied by the growing behavior (Zou et al., 2013, Corwin et al., 2016b). Furthermore, visible lesion area is correlated with the fraction of fungal RNA across lesion development stage (Windram et al., 2012) and across the 98 Botrytis strains on multiple host-genotypes (Zhang et al., 2019). Thus, lesion area is a useful approximation of the interaction between the fungal and plant genetic variation.

**Fig. 3:**
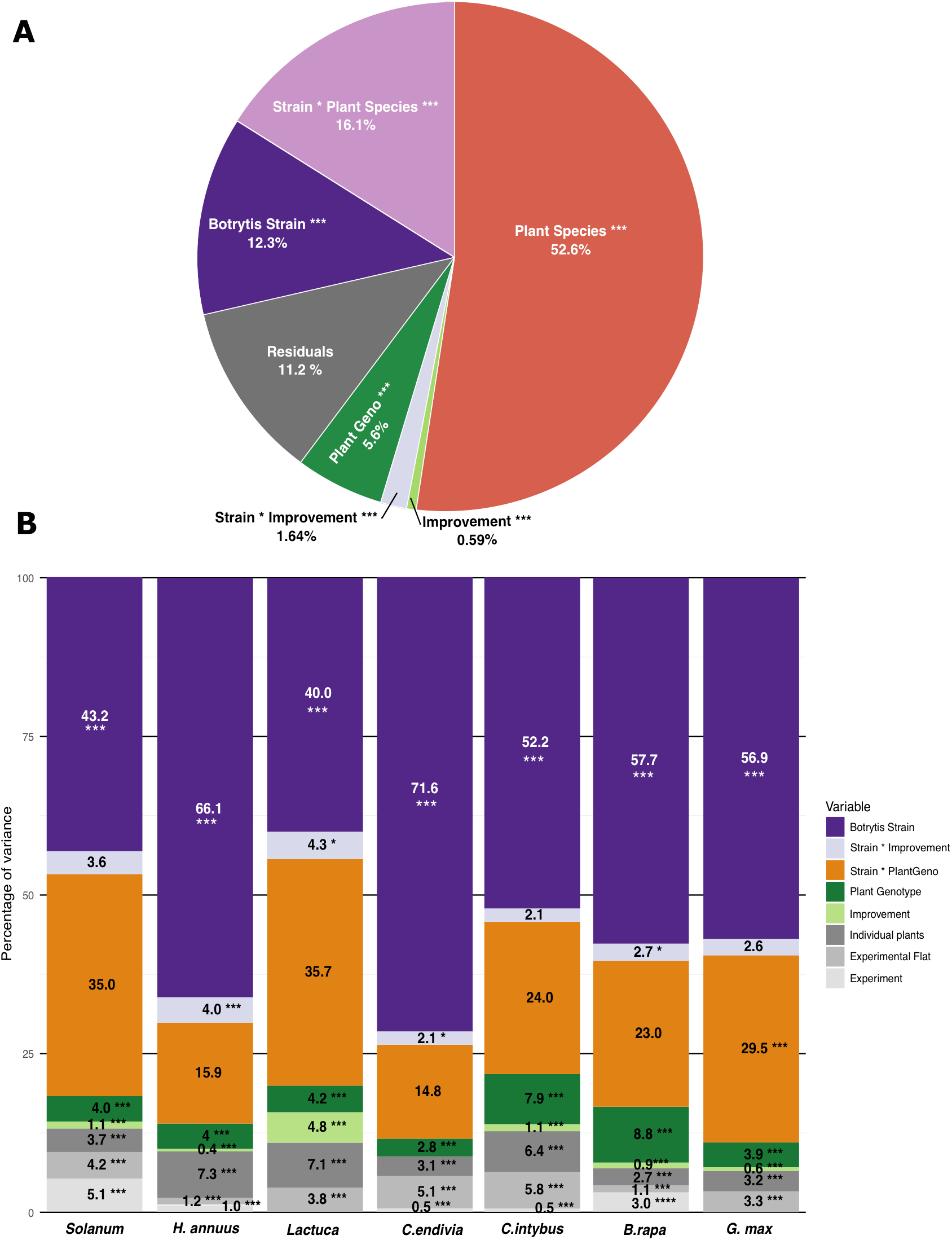
Host variation predominates the outcomes of plant-Botrytis interactions across species while Botrytis variation predominates within plant species. A) Multi-host linear model estimating the contribution of plant species, plant genotypes, level of improvement, Botrytis strains and their interaction on the percentage of variance in lesion area. B) Species-specific linear mixed models that estimate the percentage of variance in lesion area. In grey are the experimental parameters classed as random factors. Two-tailed t-test: *p<0.05, **p<0.01, ***p<0.005.

### Parsing Host Resistance and Botrytis Virulence Interactions

The Botrytis strain collection contains extensive genetic diversity with no evidence of population stratification by host-of-origin or geographical origin (Fig S3). This absence of structures is consistent whether genome-wide SNPs (Fig. S3A) or microsatellites (Fig. S3B) was used. Given the recombination in this population, it can be considered an admixed pool of virulence mechanisms with which to challenge the hosts (Table S2) (Atwell et al. 2018, Zhang et al. 2019, Soltis et al. 2019).

Mean lesion areas across the eight eudicot species challenged with the strain collection show a wide distribution, ranging from no visible damage to over 4cm^2^ of necrotic leaf tissue after 72 hours of infection (Fig. 2). *A. thaliana* has the lowest susceptibility while *H. annuus* is the most susceptible species (Fig. 2). Within each species, the effect of genetic variation between the Botrytis strains (40-71% of the explained variance) and their interaction with the host genotype (15-35% of the explained variance) controls the vast majority of the variance in the lesion area (Fig. 3B, Fig. S4B, Table S3).

Using the least-square mean lesions from each host genotype x Botrytis strain interaction (6 replicates), we extended this to a multi-host analysis across all seven host species. This multi-host analysis showed that differences in Botrytis virulence between species are the main determinant of lesion area accounting for 52% of the total variance (P < 2.2×10^−16^; Fig. 3A, Table S3). Further, the interaction between the host species and Botrytis strains (16% of total variance; P < 2.2×10^−16^) matters more than the Botrytis strains alone (12% of total variance; P < 2.2×10^−16^). Finally, the host of origin for each strain (1.7% of total variance; p< 2.2×10^−16^) and the geographical origin (0.2% of total variance; p=0.0015) have small effects on Botrytis virulence or specificity across the species tested (Fig. 4, 5), confirming the low adaptation to specific hosts or population structure in Botrytis (Atwell et al., 2018, Soltis et al., 2019). As such, the host-Botrytis interactions are additively controlled by large to narrow variations in the host genetic diversity in interaction with strain specific pathogenicity.

**Fig. 4:**
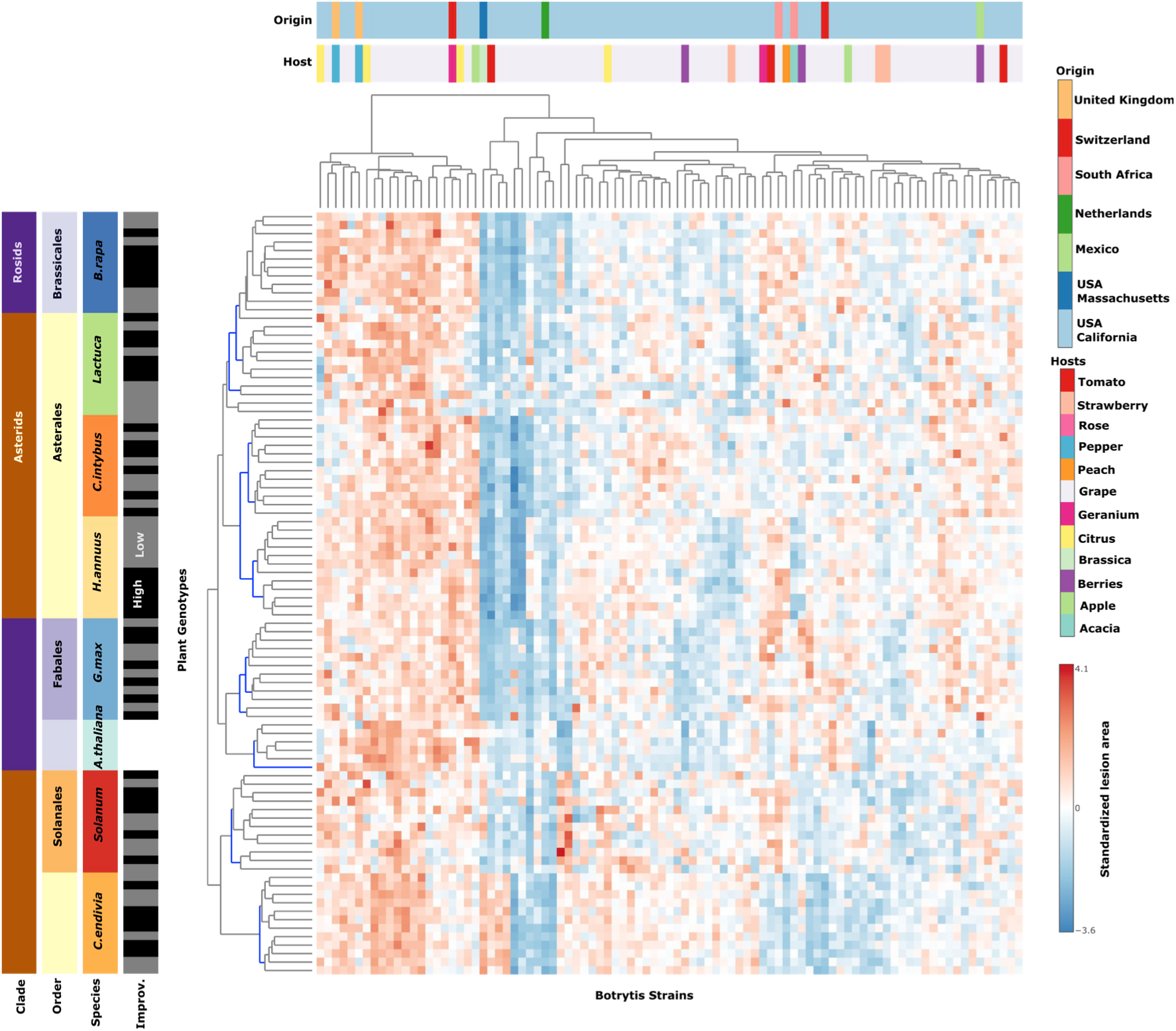
Variation of plant susceptibility in the Botrytis pathosystem does not track plant evolution. Heatmap of standardized (z-scored) least-squares means of lesion area (n=6) for Botrytis strains (x-axis) interacting with 90 plant genotypes (y-axis). The strains were isolated largely in California (light blue color in the origin bar) and on grape (light purple in the host bar). For *A. thaliana*, five single gene knockout mutants and the corresponding wild-type Col-0 are presented. For the seven crop species, six genotypes with low (grey) and six with high (black) level of improvement are presented. The seven crop species were chosen to represent a wide spectrum of phylogenetic distances across Rosids (Brassicales and Fabales) and Asterids (Asterales and Solanales). Branches in the dendrogram that are supported with 95% certainty after bootstrapping are indicated in blue. No branches in the Botrytis strain dendrogram were significant.

**Fig. 5:**
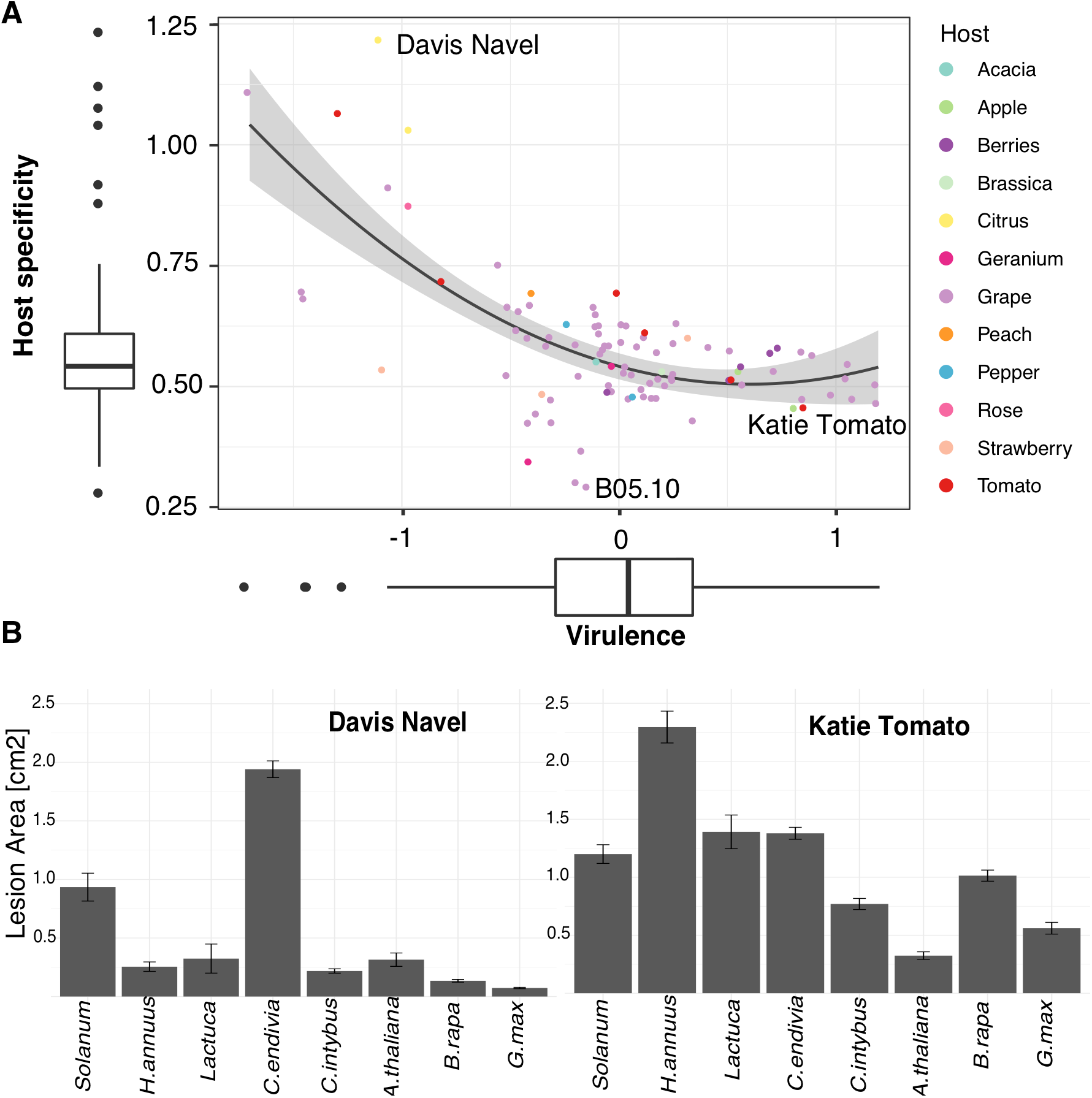
In the Botrytis strain collection, low virulence is linked to high host specificity. A) Estimates of general virulence and host specificity across eight eudicots for the Botrytis strains are shown. The strains are colored according to the plant host from which they were collected. A quadratic relationship was the optimal description of the relationships between specificity and virulence and is shown with a grey confidence interval (R^2^=0.44, P=6.36×10^−13^). B) Mean lesion area and standard error (n=12, except *A. thaliana* n=6) across the eight eudicot species is provided for two strains at the extremes of the host specificity/virulence distribution.

### Botrytis and crop domestication

For each crop species, high and low improvement genotypes were included (Table S1). In addition to sampling the within species genetic diversity, this design was chosen to assess how seven distinct domestication histories targeting seed, fruit or leaf (see Material & Method) may influence susceptibility to Botrytis. Domestication in all of these species has been associated with significant changes in their interactions with insects and specialist pathogens (Whitehead et al., 2017) (Karasov et al., 2014, Whitehead et al., 2017, Stukenbrock and McDonald, 2008, De Gracia et al., 2015). While, none of the crops species have been explicitly bred for resistance to Botrytis, domestication of lettuce and chicory did shift their growing conditions towards leaves densely compacted and adaptation to colder and wetter environments, two traits associated with increased Botrytis prevalence (De Vries, 1997, Mou, 2011, Dempewolf et al., 2008).

When infected with Botrytis, tomato and lettuce showed a decrease in resistance in high improvement genotypes (Fig. 2). The other five species (sunflower, endive, chicory, *Brassica* and soybean) showed an increase of Botrytis with crop improvement (Fig.2). While the effect of crop improvement on lesion area is statistically significant (P< 2.2×10^−16^), this effect is exceedingly small, 0.6% of the total variance across all plant species and 2-4% of variance within specific species models (Fig. 3, Table S3). Further, the range of variation in resistance to the diversity of Botrytis strains was similar between wild and domesticated lines for all species. This bi-directional small effect of domestication on this generalist pathogen is likely an indirect effect of other selective pressures in domestication and illustrates the challenge of predicting resistance to a generalist pathogen (Soltis et al., 2019).

### Botrytis and the Eudicots

To assess how the evolutionary relatedness of defense components may affect the Botrytis interaction, we utilized the standardized least-square mean lesion area of every Botrytis strain on every plant. Unsupervised clustering showed that variation in host-Botrytis interactions identifies genotype groupings for all species, i.e. the virulence of the Botrytis strains across the host genotypes is most similar within a host species. However, beyond the individual species level, the relatedness between hosts did not correlate with the relatedness between the measured host-pathogen interactions (Fig. 4). For example, phylogenetically distant crops such as *B. rapa* and *Lactuca* have a similar susceptibility pattern to the 98 Botrytis strains. In contrast, the two sister species, *C. endivia* and *C. intybus*, have highly divergent susceptibility patterns. Thus, the interaction of Botrytis with host plants is predominantly defined by variation at the narrow species-by-species level with minimal comparability between species of a particular family or genus.

To test whether these species-by-species interactions with Botrytis extend to genotypes with single alterations in plant defense, we included data for *Arabidopsis thaliana* Col-0 and five single gene knockout mutants (*coi1*, *anac055*, *npr1*, *tga3*, *pad3*) *(Rowe et al., 2010, Zhang et al., 2017)*. These mutants have a larger effect on plant susceptibility (39% of the variance in lesion area; Fig. S4, Table S3) in comparison to variation between crop genotypes (4-8.8% of the variance within a species; Fig. 3B). While these mutants abolish major sectors of plant resistance, the Botrytis-host interactions still cluster these mutants to wild-type Arabidopsis (Fig. 4) with branch lengths comparable to variation between genotypes within crop species. Overall, Botrytis disease symptoms reveal the complexity of host-specific interactions within the eudicots.

### Botrytis balances host specificity and virulence

The pressure to specialize on fungal pathogen varies across the continuum of lifestyles. While biotrophs are under strong directional pressure to develop specific and intricate responses to their host, saprophytes that eat decaying plant tissues have low pressure to specialize (Barrett et al., 2009, Krah et al., 2018, Leggett et al., 2013). For a generalist necrotroph as Botrytis, the level of host specificity of individual strains is largely unknown.

Host specificity is usually calculated qualitatively based on the broadness of presence/absence of symptoms on hosts. As Botrytis is a polyphagous necrotroph with quantitative disease symptoms, we estimated host specificity as the variation in disease symptom across hosts. With such estimates, generalist strains will tend to have equal virulence on various host, while more specialized strains will have increased virulence on a host relative to the other hosts. To account for variations in resistance among hosts, we used the coefficient of variation of least-square mean of lesion area across the eudicot species (Fig. 5B). Correspondingly, we estimated the general virulence for each Botrytis strain as the mean of standardized lesion area across the eudicot species (Fig.5). This creates a dataset to compare host specificity and overall virulence within a generalist pathogen.

The Botrytis strains cover a range of host specificity and virulence (Fig.5). B05.10, the reference strain for the *Botrytis cinerea* genome and the strain used in >90% of all papers to assess host susceptibility to Botrytis, is the strain with the lowest host-specificity/highest generalist behavior (Fig. 5). Strains with increased host specificity had on average lower virulence across all eudicots (Fig. 5). To check whether this trend was an artifact due to a potential relation between standard deviation and mean of lesion area, we investigated host specificity as the lesion area of each strains optimal host versus the strains average lesion area on the other 7 hosts (Fig. S5). Each strains deviation from theoretical generalism (equal virulence on all hosts, Fig. S5B) revealed a quadratic relation matching the relation between coefficient of variation based host specificity and virulence (Fig. 5). Both metrics revealed that strains with increased host-specificity were rare in the population. In combination, this suggests that Botrytis is under pressure to maintain broad host ranges and moderate virulence within individual strains.

### The genetic architecture behind Botrytis interactions

Various genetic architectures can explain the specificity and virulence of pathogens. Pathogens such as *Fusarium oxysporum* and *Alternaria alternata* comprise individual strains that display host specialization linked to disposable chromosomes (van der Does and Rep, 2007, Bertazzoni et al., 2018, Mathee et al., 2008). Other fungal plant pathogens such as *Ustilago maydis* have compartmentalized regions with rapidly evolving gene clusters (Moller and Stukenbrock, 2017).

To estimate the respective genetic architecture of virulence and host specificity in Botrytis, we conducted a genome-wide association study (GWAS). GWAS revealed that both overall virulence across eight eudicots and host specificity are highly polygenic traits with respectively 4351 and 5705 significant SNPs at a conservative 99.9% threshold (Fig. 6, 7A).

**Fig. 6:**
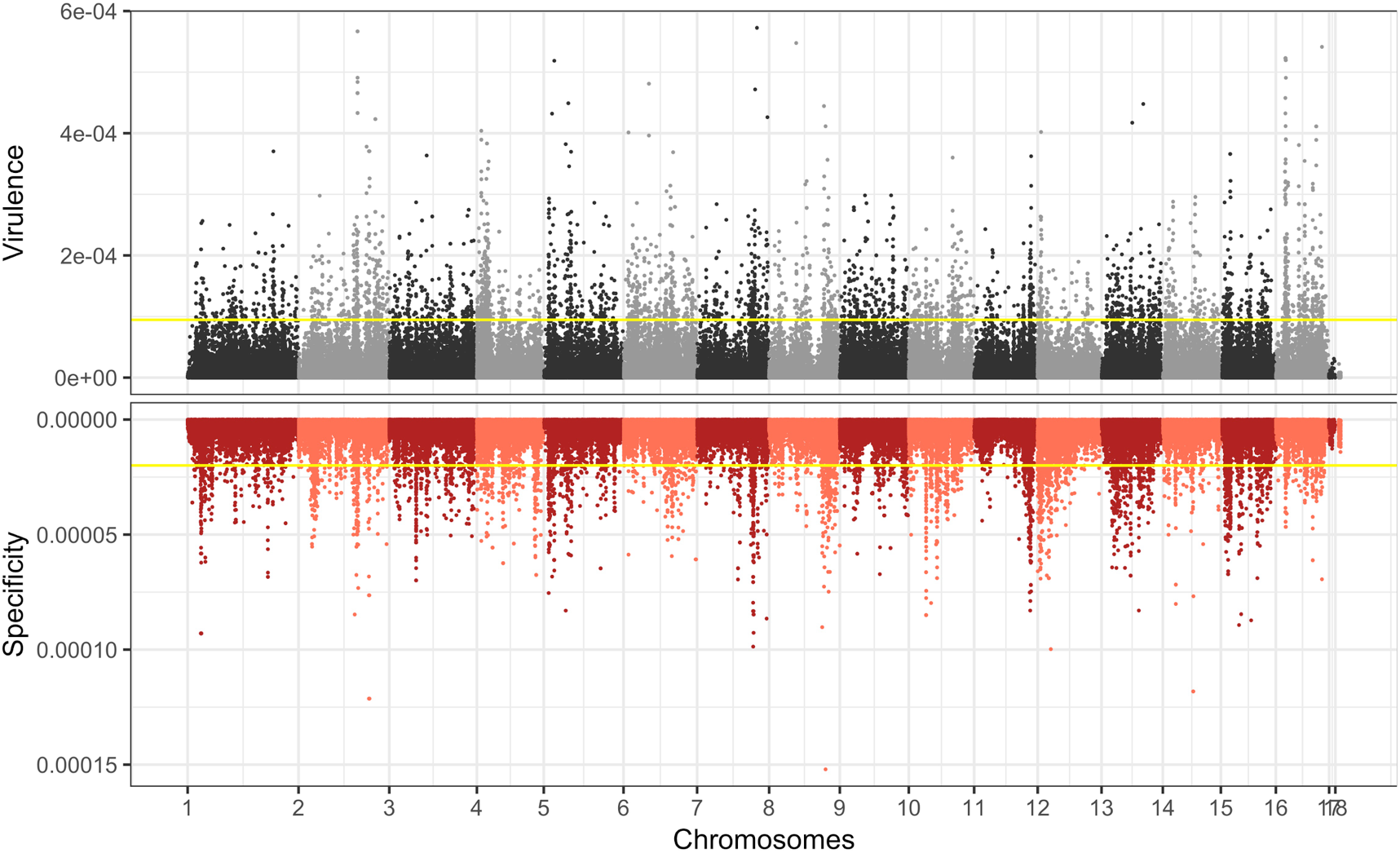
Botrytis virulence and host specificity genetic architecture is polygenic. Effect size of 271,749 SNPs with reference to B05.10 genome estimated through ridge regression GWAS. In grey is plotted the effect on general virulence and in red is the effect on host specificity. The yellow lines represent the conservative significance threshold at 99.9% for each trait as determined by permutation.

**Fig. 7:**
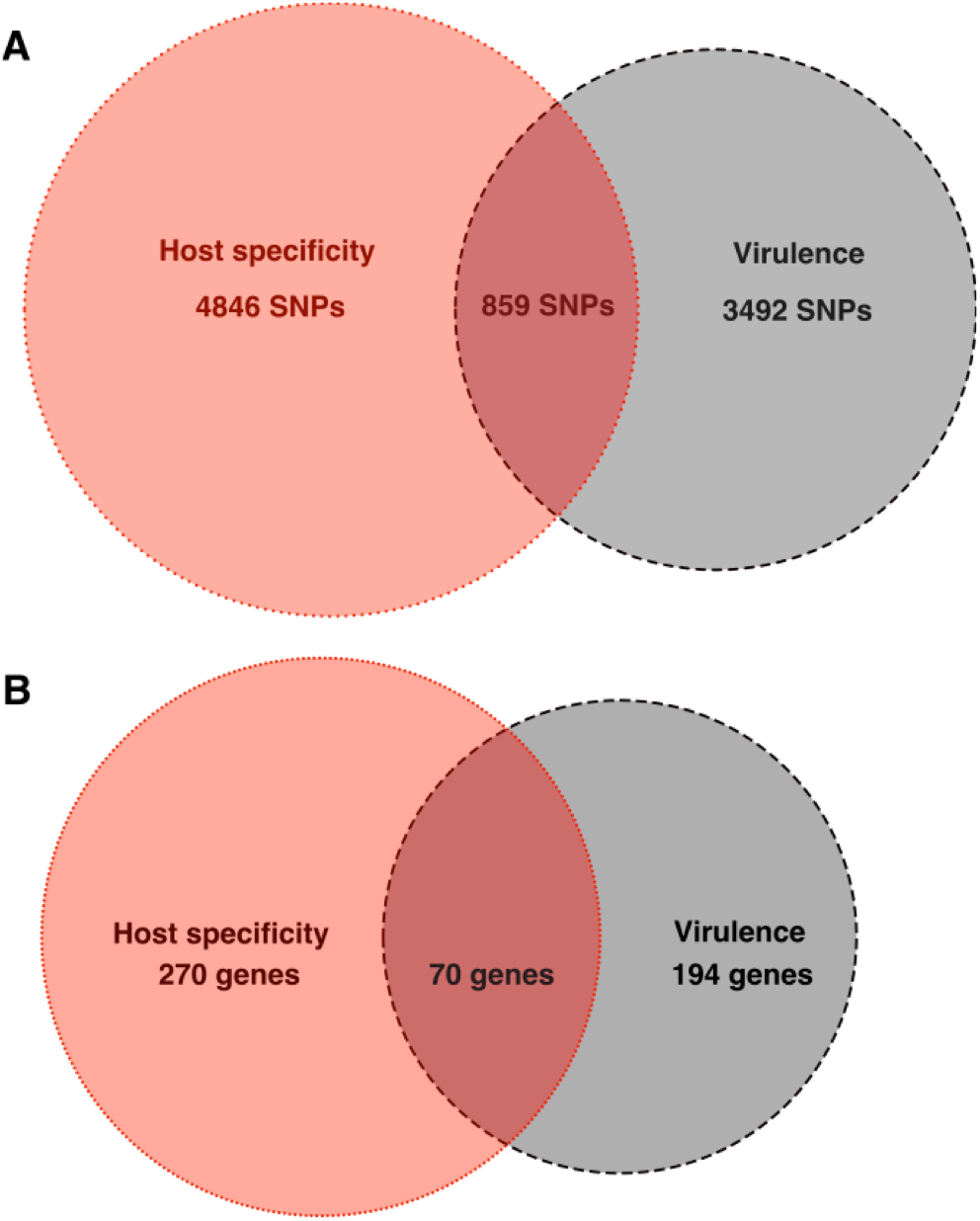
GWAS show that host specificity and virulence are different traits. A) Significant SNPs associated to host specificity, virulence or both. B) Genes with SNPs potentially affecting the function of the protein associated to host specificity, virulence or both.

By using the effect sizes of SNPs significant at the 99.9% threshold to create a genomic prediction vector, we show that the associated SNPs capture a significant fraction of the variance in the Botrytis strains virulence (R^2^=0.74, P <2.2×10^−16^, Fig. S6A) and host specificity (R^2^=0.63, P <2.2×10^−16^; Fig. S6B). The significant SNPs are spread across 16 of the 18 chromosomes and are of small effects. None of the SNPs on chromosome 17 and 18, hypothesized to be potential disposable chromosomes (Bertazzoni et al., 2018), were associated with virulence or host specificity in this dataset (Fig. 6). This suggests that in contrast to other pathogens, Botrytis doesn’t have specific genomic structures for host specificity and virulence. Filtering for SNPs annotated as having a high functional effect revealed 194 genes that might be associated with virulence and 270 genes might be associated with host specificity (Fig. 7B, Table S4).

This GWAS reveal that virulence and host specificity are two separated traits as only 9% of significant SNPs and 70 genes with functional mutations were significantly associated to both traits (Fig 7, Table S4).

## Discussion

Pathogens form a continuum of behavior from specialists that are dependent on limited number of host species to generalists that range from conglomerates of specialized strains to polyphagous species. This study suggests that *Botrytis cinerea* may be at the end of the generalist continuum, a pathogen harboring large genetic diversity, sensitive to the genetic diversity of its various hosts and with a possible advantage to remain generalist.

### Botrytis strains are preferentially generalist with moderate virulence

By generating an infectivity matrix with a collection of 98 strains, we show that *Botrytis cinerea* contains the genetic diversity to display a wide range of virulence on various hosts. The vast majority of *Botrytis cinerea* strains are generalists with only a few strains showing a propensity to prefer an individual host among the species tested. Generalist strains may have an advantage as they maintain moderate virulence on a large spectrum of hosts. Specialization may occur via an increase in virulence on a specific host at the cost of virulence on other hosts. This specialization syndrome is illustrated by Botrytis strains with high virulence on *C. endivia* (Fig. 4), clustering because of their lack of virulence on all the other hosts rather than their increase of virulence on this host. This is different from the usual specialization models that predict that specialized strains arise from a population of weakly virulent generalists (Woolhouse et al., 2001,Barrett et al., 2009, Leggett et al., 2013), with specialization presenting an advantage. This strategy in Botrytis might be favoured by host jumps at high frequency and low speciation rate (Navaud et al., 2018).

The genetic architecture of *Botrytis cinerea* is different than other multi-host pathogens. Instead of defined genomic regions controlling host specificity, the moderate virulence of generalist strains and Botrytis ability to overcome ecological challenges might derive from the extensive genome-wide standing variation within populations (Atwell et al., 2015a, Atwell et al., 2018, Walker et al., 2017, Woolhouse et al., 2001).

When coupled with recombination, the standing variation can generate quickly diverse genetic responses (McDonald and Linde, 2002, Woolhouse et al., 2001). In support of this hypothesis, Botrytis strains in the field showed extensive recombination occurring in a seasonal pattern (Fillinger and Elad, 2015, Walker et al., 2014). Additionally, Botrytis virulence and specificity may also depend on structural variation and transposable element dynamic (Amselem et al., 2011, Kecskeméti et al., 2014). Structural variation produces variation in production of toxic metabolites (such as botcinic acid and cyclic peptides) that affect the virulence (Zhang et al., 2019, Valero-Jimenez et al., 2019). Transposons were associated to the production of diverse small RNAs released to silence host defenses (Weiberg et al., 2013). While more work is required to assess the worldwide evolution of virulence and host specificity in Botrytis including transposons and structural variations, the relative lack of strains with any host-preference in our collection does suggest that this pathogen may be under pressure to maintain a generalist life style.

### An additive quantitative model for plant pathogen interactions

The absence of phylogenetic signal between host species and the small effect of narrow lineage genetic variations may be explained by the quantitative and polygenic genetic architecture of plant resistance. While plant resistance is constituted as multi-layer systems with components of small to moderate effects (Corwin et al., 2016, Corwin and Kliebenstein, 2017, Soltis et al., 2019), the summation of these quantitative effects is enough to generate diverse resistance responses. These responses change even among closely related species, such as *C. endivia* and *C. intybus* but not among domesticated species such as in lettuce and tomato. Furthermore, the quantitative and polygenic architecture of plant resistance mirror the genetic architecture for host specificity and virulence in the pathogen. The disease outcome is therefore complex, possibly resulting of the interaction of the host and pathogen genome-wide variations.

Botrytis virulence and specificity patterns as described in this study are at the opposite of the patterns of plant-pathogen interactions generally described by co-evolutionary arms-race scenarios. In the arms-race model, defense innovations in the host select counter mechanisms in the pathogen driving by evolution of large effect genes linked to specific interactions. As such the arm-race model show strong directional pressure in pathogens to become specialists (Barrett is expected to increase with the evolutionary distances (Gilbert and Parker, 2016, Schulze-Lefert and Panstruga, 2011, Gilbert and Webb, 2007). Other qualitative models such as the inversed gene-for-gene, matching alleles or inversed matching alleles (Morris and Moury, 2019) don’t fit *Botrytis cinerea* well either. Within an additive quantitative model, Botrytis genetic diversity may vary through frequent opportunistic host switches rather than co-evolution with multiple hosts. More research is needed to verify and validate models on the evolution of pathogenicity in *Botrytis cinerea*.

## Methods

### Plant material and growth condition

Eight species were chosen to represent a wide diversity of phylogenetic distance, geographical origin and histories across the core Eudicot (Fig. 1). For simplicity in the language, we refer to those as ‘species’ although referring to taxa would be more appropriate for tomato and lettuce for which we selected sister species for wild and domesticated genotypes. When referring to *Lactuca* (lettuce), *Lactuca sativa* was sampled for improved varieties and *Lactuca serriola* for wild accessions (Walley et al., 2017). When referring to *Solanum* (tomato), *Solanum lycopersicum* was sampled for improved varieties and *Solanum pimpenellifolium* for wild accessions native from Ecuador and Peru (Lin et al., 2014).

This assay considers four Asterales species/taxa (*Helianthus annuus*, *Cichorium intybus*, *Cichorium endivia*, *Lactuca*), one Solanales (*Solanum*), one Fabales (*Glycine max*) and two Brassicales (*Brassica rapa, Arabidopsis thaliana*) (Fig. 1, Fig. 3, Table S1). *Arabidopsis thaliana* data was used as a reference for genotypes with altered plant susceptibility (Fig. S4). This reference dataset is composed of Col-0 and five knockout mutants altering plant immunity (Zhang et al., 2017, Atwell et al., 2018), through the jasmonic pathway (anac055, coi1), salicylic pathway (npr1, tga3) and camalexin pathway (pad3), an anti-fungal defense compound in *Arabidopsis*.

While the chosen species partially represent the eudicot phylogeny (Fig. 1), they were also chosen to represent the diversity of plant domestication syndromes and geographical origins (Meyer et al., 2012) (Fig. S2). *H. annuus* and *G. max* were domesticated for seeds, *Solanum* for fruit (Lin et al., 2014) while *Lactuca*, *C. intybus* and *C. endivia* were domesticated for leaf and root (Dempewolf et al., 2008). All of these species underwent a single domestication event while *B. rapa* domestication is more complex (Bird et al., 2017). *B. rapa* was domesticated and re-selected multiple times for multiple traits including seed, leaf and root characteristics. For each species, twelve genotypes were selected, including six genotypes with low (wild or landrace) and six with high (cultivar, inbred lines) level of improvement (Table S1). The genotypes were chosen based on description of domestication status, and phylogeny for each species (Blackman et al., 2011, Walley et al., 2017, Lin et al., 2014, Dempewolf et al., 2008, Bird et al., 2017, Liang et al., 2014, Valliyodan et al., 2016). Landrace genotypes were selected for *C. endivia* for which no wild relative is known and two genotypes of *B. rapa* (Fig. S2, Table S1). For soybean the comparison was within *G. max* as the growth behavior, vining, and growth conditions, tropical, for wild soybean, *G. soja*, was sufficiently different as to unnecessarily confound the comparison.

*C. endivia*, *C. intybus*, *B. rapa*, *G. max and A. thaliana* seeds were directly sowed in standard potting soil. *Solanum* and *Lactuca* seeds were bleach-sterilized and germinated on wet paper in the growth chamber using flats covered with humidity domes. After 7 days, the seedlings were transferred to soil. Seed surface sterilization and scarification was used for *H. annuus* to increase seed germination. Seeds were surface sterilized in 30% bleach for 12 minutes, followed by rinsing with sterilized distilled water, and then soaked in sterilized water for 3 hrs. ¼ of the seeds were cut off from the cotyledon end, then placed in 100 mg/L Gibberellic acid for 1 hour, followed by rinsing several times with sterilized distilled water. Treated seeds were then put in covered sterilized Petri dishes with wet sterilized germination disks at 4°C for 2 weeks and sowed.

All plants were grown in growth chambers in pots containing Sunshine Mix#1 (Sun gro Horticulture, Agawam, MA) horticulture soil at 20°C with 16h hours photoperiod at 100-120 mE light intensity. All plants were watered every two days with deionized water for the first two weeks and then with nutrient solution (0.5% N-P-K fertilizer in a 2-1-2 ratio; Grow More 4-18-38). All infection experiments were conducted on mature (non-juvenile) fully developed leaves collected on adult plants that grew in these conditions for four to eight weeks (Table S5) to account for the different developmental rates. As it is challenging to fully compare developmental stages across species (soybean stages are defined by nodes, sunflower by leaf size and so on), all leaves for the assays were collected on plants in the vegetative phase at least several weeks before bolting initiation to minimize ontogenetic effects.

### Botrytis collection and growth condition

This study is based on a collection of 98 strains of *Botrytis cinerea.* The collection samples the *B. cinerea* strain diversity across fourteen plant hosts and in smaller degree across geographical origins. Ninety percent of the strains were isolated in California, largely in vineyards (70% of strains were collected on grape), while the remaining 10% of the collection are worldwide strains (Table S2). The vineyards were in close proximity (<0.25 miles) to production and research fields growing lettuce and tomato. Further, the vineyards were not maintained and had a large number of native and invasive Asterales and Brassicales weeds. Thus, any obtained isolate is likely to have been exposed to a broad range or hosts throughout the year. The strain collection contains a large level of genetic diversity and there is no significant stratification of genetic diversity by either geographic or host species origin (Atwell et al., 201a, Atwell et al., 2018). Given the polygenic nature of virulence in Botrytis, if these isolates had adapted specifically to growth on grapes, there would have been stratification between the grape and non-grape isolates. The spore collection is maintained for long-term preservation as conidial suspension in 60% glycerol at −80°C. The strains were grown for each experiment from spores on peach at 21°C for two weeks.

### Detached leaf assay

To maximize the comparability across such a diverse collection of wild relatives and crop species domesticated for different trait, leaves were chosen as a common plant organ. Detached leaf assays were conducted following (Corwin et al., 2016, Denby et al., 2004). The detached leaf assay methodology has been used for testing plant susceptibility to plant pathogens in more than 500 scientific publications and is considered a robust method when coupled with image analysis.

In brief, leaves were cut and added to trays in which 1cm of 1% phyto-agar was poured. The phyto-agar provided water and nutrients to the leaf that maintained physiological functions during the course of the experiment. Botrytis spores were extracted in sterile water, counted with a hemacytometer and sequentially diluted with 50% grape juice to 10spores/ul. Drops of 4ul (40 spores of Botrytis) were used to inoculate the leaves. From spore collection to inoculation, Botrytis spores were maintained on ice to stop spore germination. Spores were maintained under agitation while inoculating leaves to keep the spore density homogeneous and decrease technical error. The inoculated leaves were maintained under humidity domes under constant light. To render a project of this size possible, a single time point was chosen to measure the plant-Botrytis interaction. Lesion area at 72 hours post infection (hpi) was chosen because the lesions are well defined on all targeted species but have not reached the edges of the leaves and thus are not tissue limited. Spore germination assays in 50% grape juice showed that strains germinate within the first 12 hours. On all species, the same pattern of growth was observed: after 48h most strains are visible within the leaf with the beginning of lesion formation. From 36h onward, Botrytis growth is largely linear and grows until the entire leaf is consumed (Rowe et al., 2010). The experiments included three replicates for each strain x plant genotype in a randomized complete block design and were repeated over two experiments, for a total of six replicates per strain x plant genotype.

### Image analysis

The images analysis was performed with an R script as described in (Fordyce et al., 2018). Images were transformed into hue/saturation/value (hsv) color space and threshold accounting for leaves color and intensities were defined for each species. Masks marking the leaves and lesions were created by the script and further confirmed manually. The lesions were measured by counting the number of pixels of the original pictures within the area covered by the lesion mask. The numbers of pixels were converted into centimeter using a reference scale within each image.

### Data quality control

A dataset of 51,920 lesions was generated in this project but not all leaves inoculated with Botrytis developed a visible lesion at 72hpi. These ‘failed lesions’ can be explained either by technical or biological failures. Technical failures, such as failed spore germination, stochastic noise or other non-virulence related issues can bias the true estimates of the mean. To partition biological from technical failures, the lesion area distribution was analyzed for each species and empirical thresholds were fixed (Table S6). A lesion below that threshold was considered a technical error only if the median of lesion area for a plant genotype - strain pair was larger than the threshold. The rationale is the following: when most lesions are of small size, the likelihood of biological reasons for such small lesion areas is high, while when the majority of lesion areas are large, the likelihood of technical error is high. 6,395 lesions (13% of all lesions) were considered as technical failures and removed from the dataset. The statistical analyses and modeling were run on both original and filtered datasets. The removal of technical failures does not impact the effect size of the estimates but their significance and allowed for more variance to be attributed to biological terms in the model and less in the random error term. This is as expected if we are partitioning out predominantly technical failures.

### Statistical analysis

All data handling and statistical analyses were conducted in R. Lesion area was modeled independently for each species using the linear mixed model with lme4 (Bates et al., 2014):

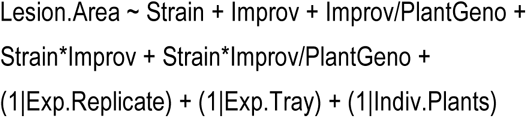

Where plant genotypes are nested within improvement level. Experimental replicate and trays as well as the individuals plants on which were collected the leaves for the detached leaf assay are considered as random factors. Plant genotypes and Botrytis strains were coded as fixed factors, although they represent random sampling of the plant and fungal species. This simplification of the model was done because previous research (Corwin et al., 2016, Fordyce et al., 2018) showed that this does not affect the effect size or significance of the estimates, while increasing dramatically the computational load. For each plant genotype, model corrected least-square means (LS-means) of lesion area were calculated for each strain from with Emmeans with the Satterwaite approximation (Lenth, 2018):

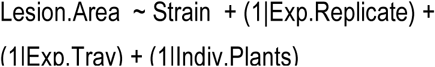

The meta-analysis model was run over all species LS-means with the linear model:

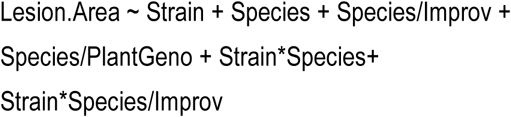

Improvement levels and plant genotypes where nested within species to account for their common evolutionary history and possibly shared resistance traits of genotypes with low and high levels of improvement. Variance estimates were converted into percentage of total variance to ease the comparison of the different models.

To visualize the relationship between strains and plant genotypes, a clustered heatmap was constructed on standardized LS-means with iheatmapr (Schep and Kummerfeld, 2017). The LS-means were standardized (z-score) over each plant genotypes by centering the mean to zero and fixing the standard deviation to one to overcome the large variation on lesion area across species and large variation in variance linked to the lesion area (Fig 2). Species with low lesion area had also small variance while species with large lesion area presented large variance. Seven strains that were not consistently infected on all 90 genotypes were dropped, as hierarchical clustering is sensitive to missing data. The unsupervised hierarchical clustering was run with the ‘complete’ agglomerative algorithm on Euclidean distances. The significance of the dendrogram was estimated with pvclust (Suzuki and Shimodaira, 2006) over 20000 bootstraps. The significance of branches was fixed at **α** =0.95. For the plant genotypes dendrogram, branches were consistently assigned across hierarchical clustering methods (both ‘complete’ and ‘average’ algorithms were ran) and bootstrapping while in the Botrytis strains dendrogram, none of branches showed consistency. The heatmap provides a global picture of how plant genotypes interact specifically with each Botrytis strains. To estimate the global virulence of each strain, we calculated the mean of the standardized LS-means of lesion area across the eight eudicot species. The host specificity was calculated from the raw LS-means as the coefficient of variation (standard deviation corrected by the mean σ/µ) across the eight species (Poisot et al., 2012). Low host specificity indicates that strains grew consistently across the eight species, while high host specificity indicates large variation in lesion area across species (Fig. 5B), therefore preference for some species.

### Genome-wide association

All strains were previously whole-genome sequenced at on average 164-fold coverage (Atwell et al., 2018). Host specificity and virulence were mapped using 271,749 SNPs at MAF 0.20 and less than 20% of missing calls with B05.10 genome as reference. Among these 271,749 SNPs, 50.2% of SNPs are in intergenic regions, 26.8% are in coding sequences, 10.2% are in 3’ UTR, 7.5% in 5’ UTR and 5.3% are in introns.

The GWA was performed using a generalized ridge regression method for high-dimensional genomic data in bigRR (Shen et al., 2013). This method, which tests all polymorphisms within a single model, was shown to be adapted to estimate small effect polymorphism (Corwin et al., 2016, Fordyce et al., 2018). Due to the modeling of polymorphism as random factors, it imputes a heteroscedastic effect model as effect size estimates rather than p-values. Other GWA methods have been tested for mapping plant-Botrytis interactions (Corwin et al., 2016, Atwell et al., 2018) and hold comparable results to bigRR model. In particular, Gemma that accounts population structure does not perform significantly better due to the low population structure in the strain collection (Atwell et al., 2018). Furthermore, BigRR approach was chosen for its known validation rate (Corwin et al., 2016, Fordyce et al., 2018) in the pathosystem, the estimation of effect size, ability to perform permutation test on significance threshold and speed of calculation on GPU. Significance of the effect size was estimated based on 1000 permutations. The 99.9% threshold was used as a conservative threshold for SNP selection although 1000 permutations allow only an approximation of such a high threshold. The genomic predictions were calculated by multiplying the effect size by their corresponding genotype and summing the resulting values over all SNPs for each strain. The genotypes were coded in reference to B05.10 genome where 0 was referring to B05.10 allele and 1 was different from B05.10. SNPs were annotated based on their location in ASM83294v1 assembly with SnpEff (Cingolani et al., 2012) while gene annotation was extracted from the fungal genomic resources portal (fungidb.org). The gene list was filtered based on the annotation of significant SNPs identified by GWAS as potentially having functional effects, i.e. non-synonymous mutation in the coding sequence), by SnpEff.

## Acknowlegments

Seeds used in this study were provided by Laura Marek and Kathleen Reitsma at the US department of agriculture, Chris Pires, the UC Davis Tomato Genetics Resource Center, the Center for Genetic Resources Netherland (CGN) through Guy Barker and Graham Teakle. The experiments were performed with the help of Dihan Gao, Aysha Shafi, David Kelly, Matisse Madrone, Melissa Wang, Josue Vega, Aleshia Hopper and Ayesha Siddiqui.

## Funding

NSF IOS 1339125 and 1021861 to DJK.

## Author contributions

Conceptualization and supervision: DJK; Data analysis: CC; Founding acquisition: DJK, Investigation: GS, CC, NS, RG, JC; Resources: SA; Writing: CC, DJK.

## Author Information

The authors declare no competing interests. Correspondence and requests for materials should be addressed to. R codes and datasets are available on github.com/CaseysC/Eudicot_Rcode.

---

**Fig. S1:**
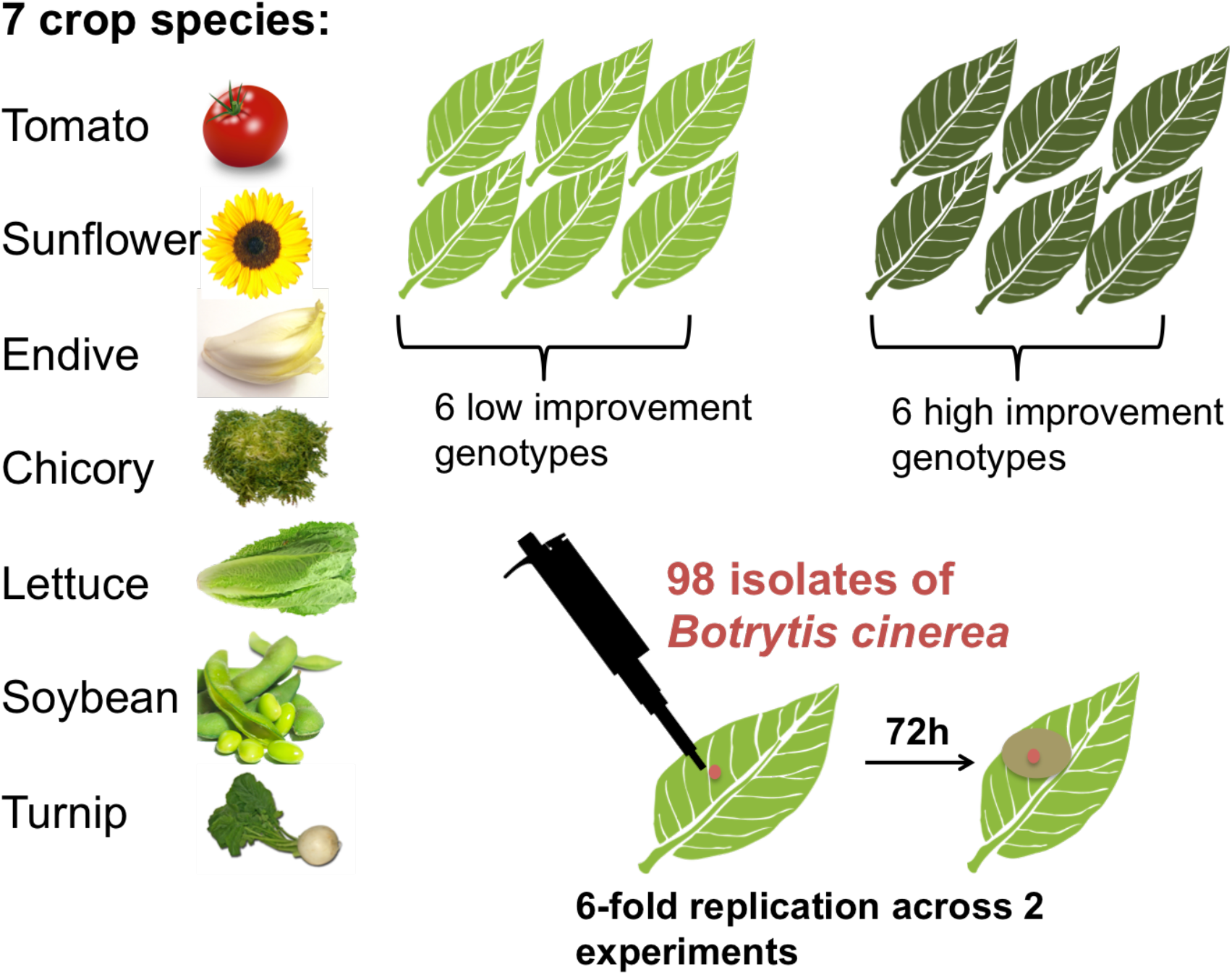
Experimental Design. We tested the virulence of 98 *Botrytis cinerea* strains on seven eudicot crop species using randomized complete block design detached leaf assays with 6 fold replication.

**Fig. S2:**
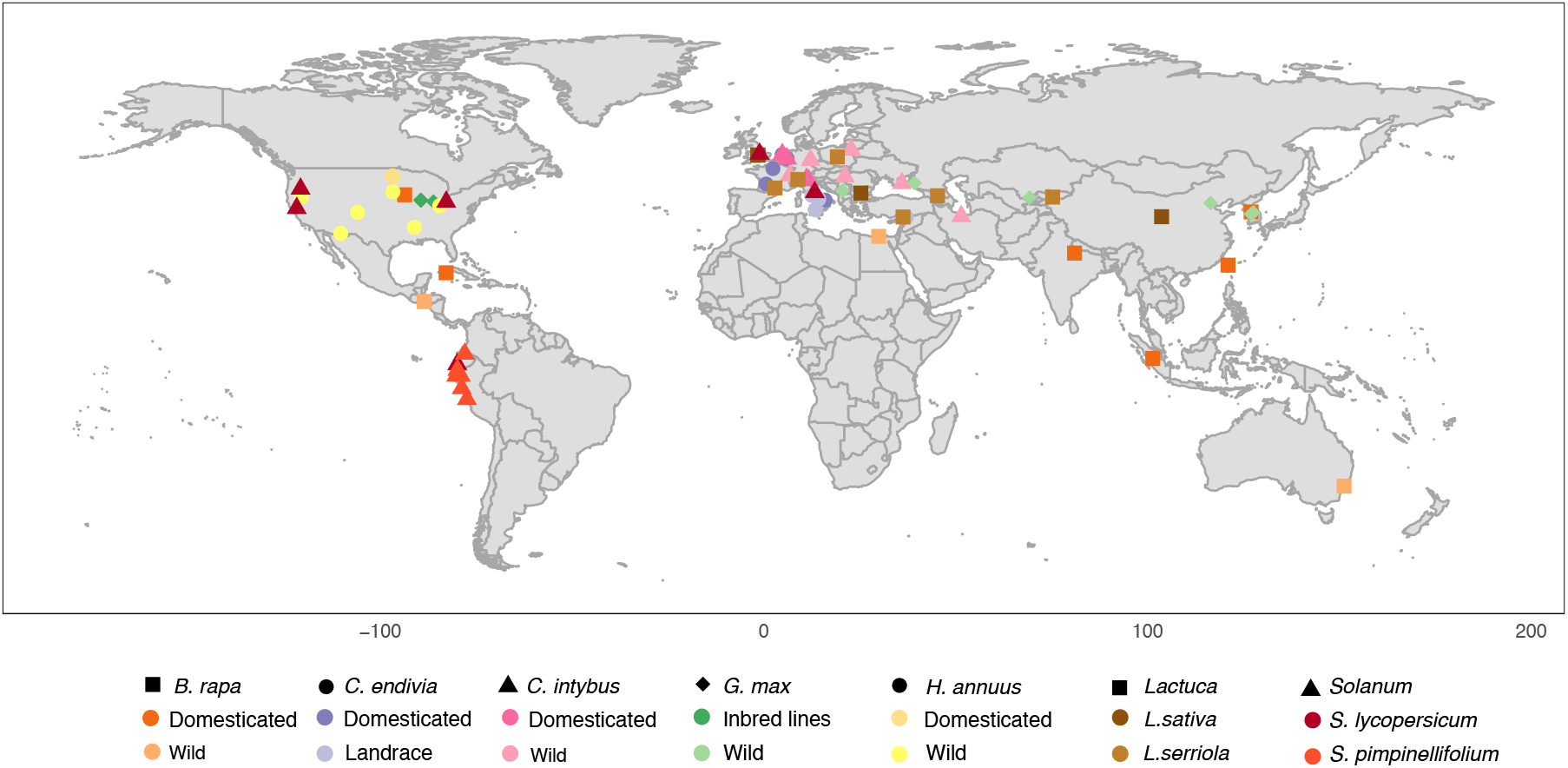
Geographical origin of the 84 Eudicot genotypes selected for this study. *B. rapa* samples come from all over the world, *C. endivia*, *C. intybus*, *Lactuca* and *G. max* come from Eurasia, *H. annuus* come from North America and *Solanum* from South America. Colored symbols represent the species and improvement status.

**Fig S3:**
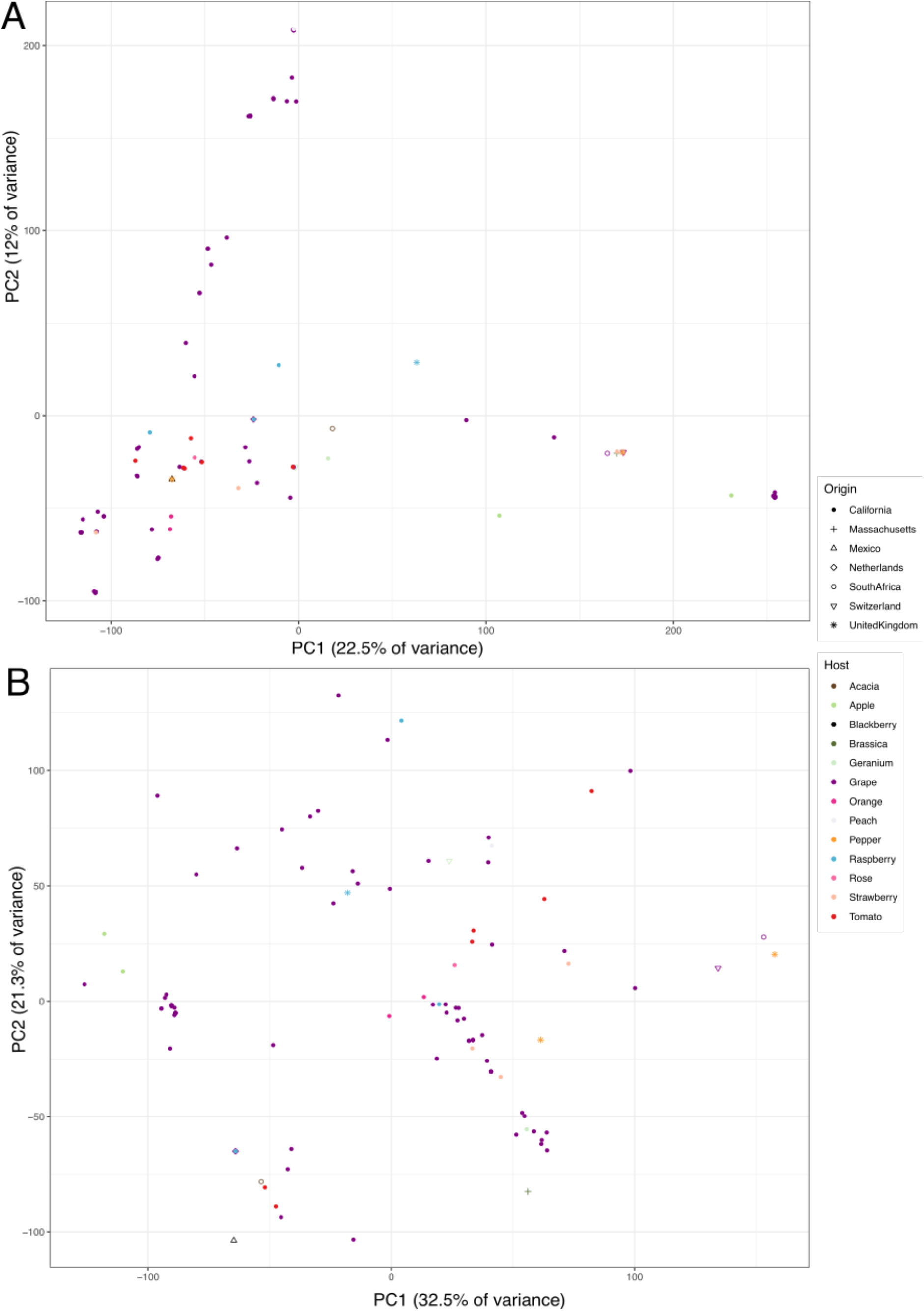
Principal component analysis of genetic variants across the 98 isolates. A) PCA based on 271749 SNPs at 20% minor allele frequency and missing data. B) PCA based on fourteen microsatellites (Diao et al., 2019). Colors represent the host of origin and the shape the geographical origin of each strain.

**Fig. S4:**
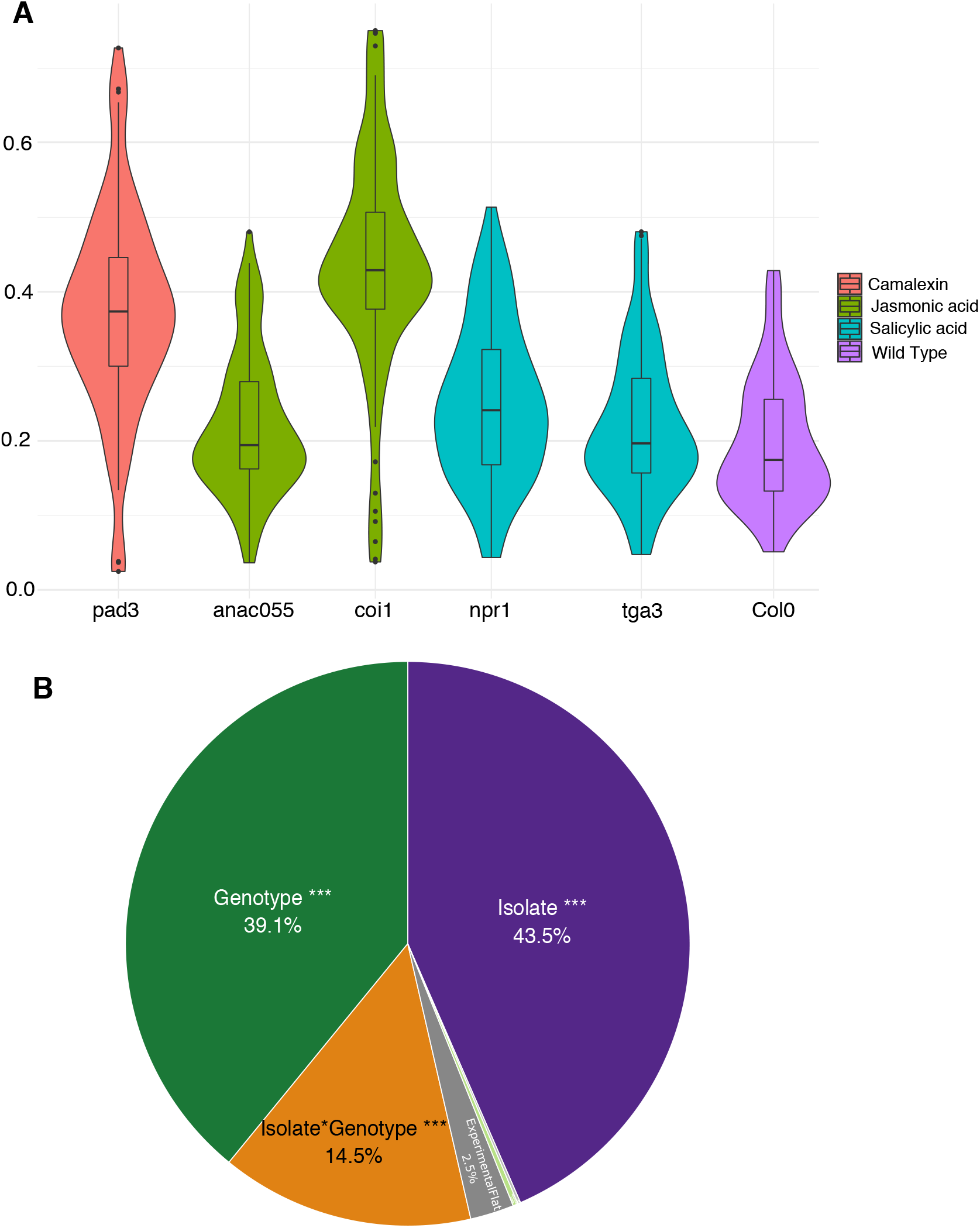
Variance in lesion area in *Arabidopsis thaliana*. A) Violin and boxplot represent the mean lesion area distribution for genotypes affecting camalexin synthesis (red), jasmonic acid signaling (green), salicylic acid signaling (blue) in comparison to the wild-type Col0 (purple) for all 98 Botrytis strains. B) Species-specific linear mixed model that estimate the percentage of variance in lesion area. In grey are the experimental parameters used as random factor. Two-tailed t-test: *p<0.05, **p<0.01, ***p<0.005.

**Fig. S5:**
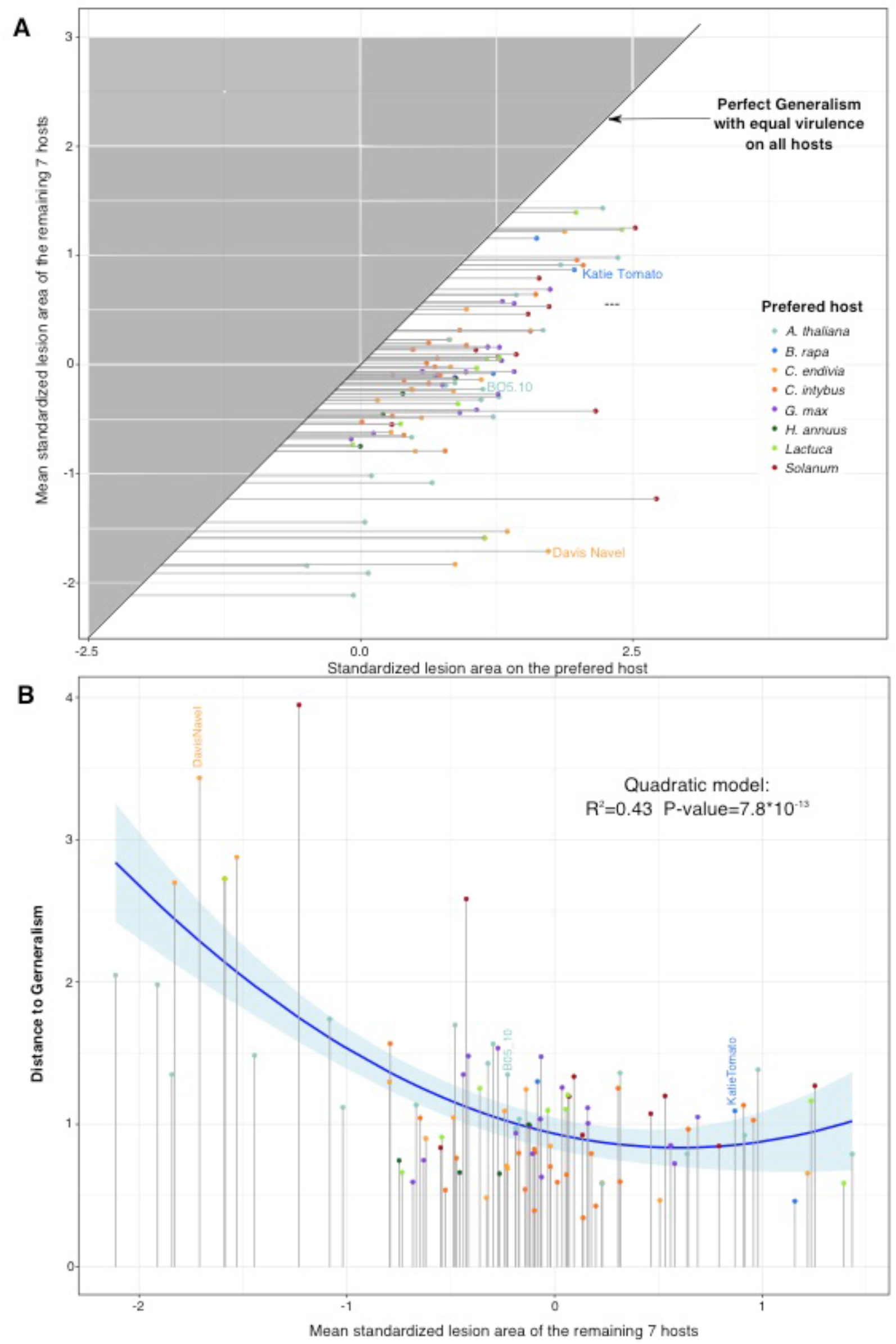
Modeling of host specificity based on mean lesion area for the 98 Botrytis strains. A) Comparison of the standardized lesion area of the preferred host (largest lesion) to the mean of the seven remaining species. The 1:1 line represent a theoretical ‘generalism’ in which strains would infect all host with equal virulence. The grey area is mathematically impossible. B) The quadratic relation between standardized mean lesion area on the remaining seven hosts and the distance to generalism confirm the trend of Figure 5, without considering the variance in lesion area.

**Fig. S6:**
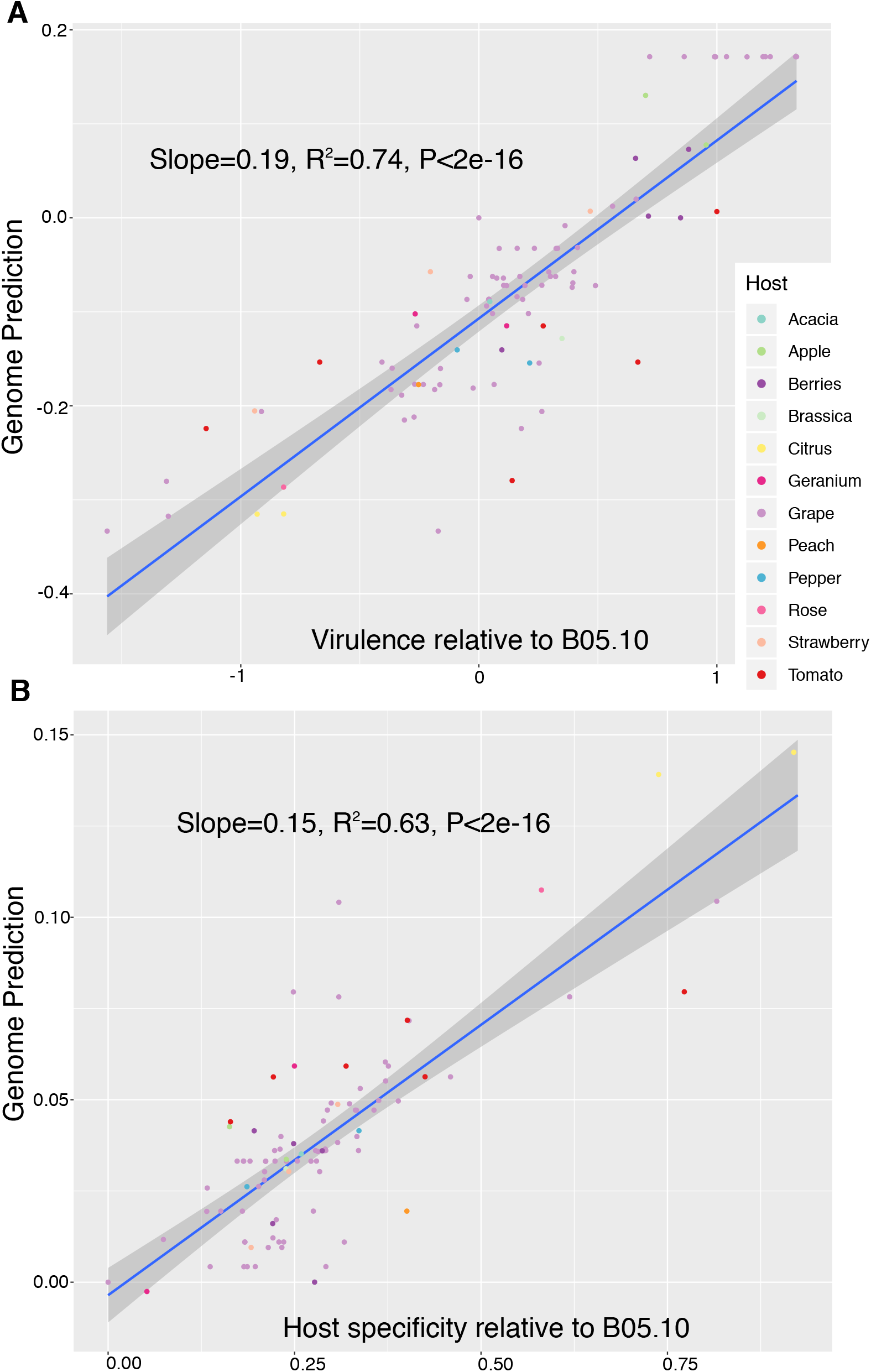
Linear model between genomic prediction based on significant SNPs at 99.9% threshold and phenotypes. A) Virulence relative to B05.10. B). Host specificity relative to B05.10. Dot colors represent the plant host Botrytis strains were collecting from. Significance was calculated with a two-tailed t-test.

## Supplementary Tables

**Table S1: Information for 84 accessions of seven crop species.** The accession ID corresponds to the germplast collections (the U.S. Department of Agriculture Germplasm Resources Information Network [USDA Grin], the Centre for Genetic resources in the Netherland [CGN], UC Davis Tomato Genetic Resource Center [TGRC]). Information on the species and subspecies, Taxon, order, clade, improvement status, cultivar name, geographical origin in addition to latitude and longitude coordinates are provided when available.

**Table S2: Information for the 98 strains of *Botrytis cinerea***. The name of the isolates, geographical origin with latitude and longitude coordinates (when available), the name and affiliation of the person who collected the strain, the year of isolation and plant species (host) on which the strain was collected are provided.

**Table S3: Detailed results for the nine linear and mixed models performed in this study, plotted in Fig. 3 and Fig. S4.** For each model, the R formula and syntax are provided. For each model and factor, the statistics (chi or sum of square), percent of variance, degree of freedom and p-values are provided.

**Table S4: Information on the 534 genes with functional effect SNPs identified by GWAS for virulence and host specificity.** Location in the B05.10 reference genome and various functional annotations are provided. An associated table provides information for each column including a description of the content, origin of the data and the reference genome of each database used.

**Table S5: Germination and growth conditions for the eight eudicot species**. All plants were grown in growth chambers at 20°C with 16h hours photoperiod at 100-120 mE light intensity for four to eight weeks to account for the different developmental rates.

**Table S6: Maximum lesion size and median thresholds.** These thresholds were used for the partitioning of technical and biological failures in developing lesions as well as the percentage of data points removed in each of the eight eudicot species.

## Notes

#### Summary of Updates

The focus of the paper was changed after concern in peer-review about the experiment design for the analysis of co-evolution dynamics. This shifts the manuscript from looking specifically at co-evolution to trying to lay out the groundwork for how genetic variation in the host and pathogen are shaping the interaction across a number of eudicot lineages. We also address reviewer comments on host specificity and deepened the GWAS analysis.

